# FDA-ARGOS: A Public Quality-Controlled Genome Database Resource for Infectious Disease Sequencing Diagnostics and Regulatory Science Research

**DOI:** 10.1101/482059

**Authors:** Heike Sichtig, Timothy Minogue, Yi Yan, Christopher Stefan, Adrienne Hall, Luke Tallon, Lisa Sadzewicz, Suvarna Nadendla, William Klimke, Eneida Hatcher, Martin Shumway, Dayanara Lebron Aldea, Jonathan Allen, Jeffrey Koehler, Tom Slezak, Stephen Lovell, Randal Schoepp, Uwe Scherf

**Affiliations:** U.S. Food and Drug Administration; U.S. Army Medical Research Institute of Infectious Diseases; Institute for Genome Sciences at the University of Maryland; National Center for Biotechnology Information, National Library of Medicine, National Institutes of Health; Lawrence Livermore National Laboratories

**Keywords:** infectious disease (ID), next generation sequencing (NGS), diagnostics (Dx), ID-NGS-Dx, agnostic (unbiased) ID sequencing, targeted ID sequencing, metagenomics shotgun sequencing, isolate shotgun sequencing, *Enterococcus avium*, Ebolavirus

## Abstract

Infectious disease next generation sequencing (ID-NGS) diagnostics are on the cusp of revolutionizing the clinical market. To facilitate this transition, FDA proactively invested in tools to support innovation of emerging technologies. FDA and collaborators established a publicly available database, FDA dAtabase for Regulatory-Grade micrObial Sequences (FDA-ARGOS), as a tool to fill reference database gaps with quality-controlled genomes. This manuscript discusses quality control metrics for the proposed FDA-ARGOS genomic resource and outlines the need for quality-controlled genome gap filling in the public domain. Here, we also present three case studies showcasing potential applications for FDA-ARGOS in infectious disease diagnostics, specifically: assay design, reference database and *in silico* sequence comparison in combination with representative microbial organism wet lab testing; a novel composite validation strategy for ID-NGS diagnostics. The use of FDA-ARGOS as an *in silico* comparator tool could reduce the burden for completing ID-NGS clinical trials. In addition, use cases identifying *Enterococcus avium* and Ebola virus (Zaire ebolavirus variant Makona) demonstrate the utility of FDA-ARGOS as a reference database for independent performance validation of new tests and for documenting how one would use this database as an *in silico* sequence target comparator tool for ID-NGS validation, respectively.

## INTRODUCTION

The Food and Drug Administration’s (FDA) premarket review of in vitro diagnostics relies on safety, efficacy, quality, and performance and ensures patient access to safe and accurate new technologies, such as next-generation sequencing. Within this premarket review, FDA performs risk-based evaluation of novel diagnostic devices by leveraging clinical expertise, as well as research evidence to support regulatory decisions and considers patient values and preferences. Infectious disease next generation sequencing (ID-NGS) diagnostics, with the potential to identify any microbial organism or genomic marker from a patient sample in a single test, are poised to enter the clinical diagnostic laboratory (Goldberg, Sichtig et al. 2015, Arnold 2017, Heger 2018). For accurate identification of any infectious organism, ID-NGS requires comprehensive reference databases, thereby strongly emphasizing the need for more complete high-quality reference genomes. Metagenomic agnostic sequencing also requires novel validation strategies as the traditional diagnostic evaluation for all known organisms is unfeasible. Described in greater detail throughout this paper (Figure 1), here we are describing one effort for defining a composite-reference method approach for ID-NGS device validation utilizing *in silico* sequence comparison.

**Figure 1:**
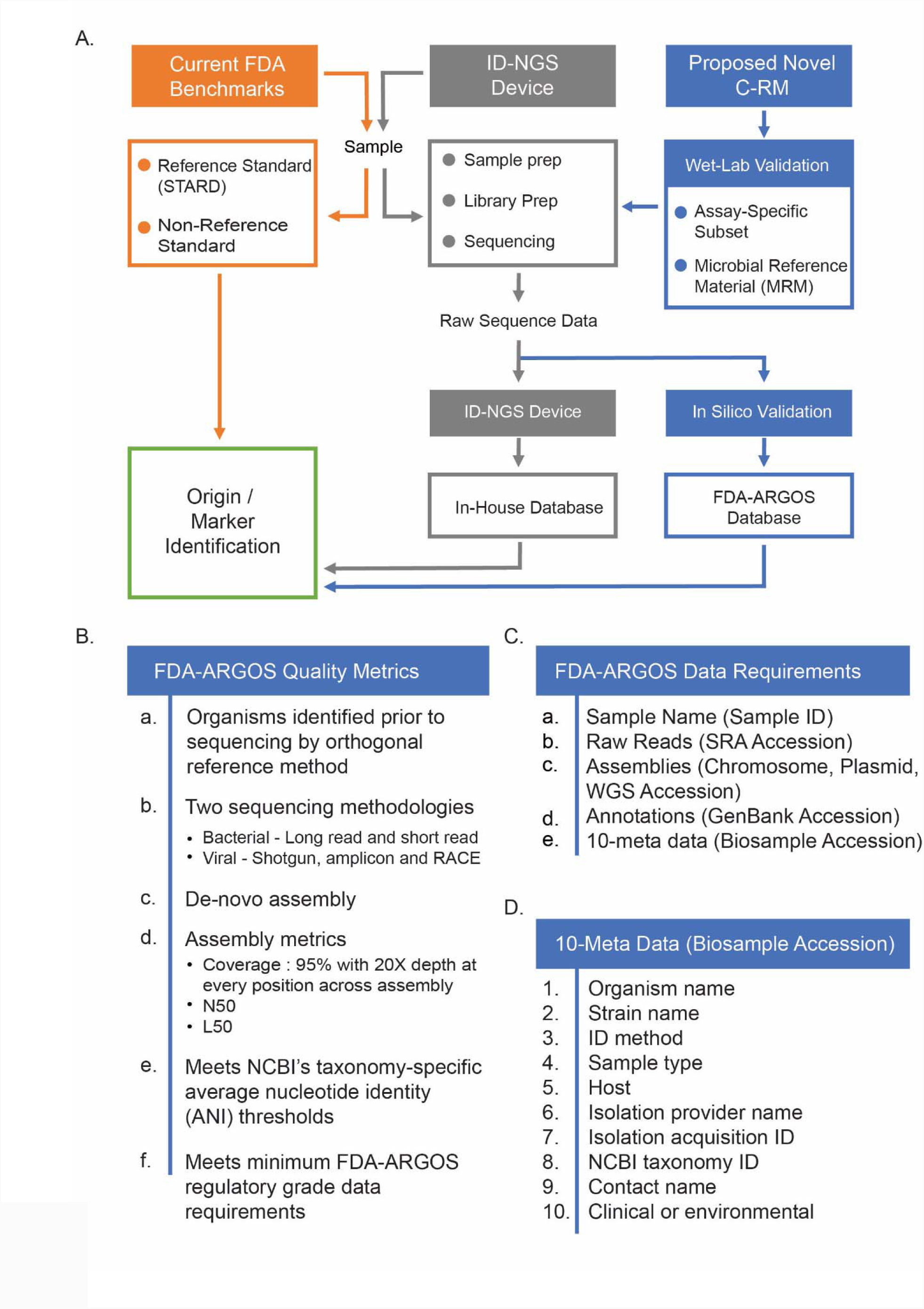
Proposed Novel Composite Reference Method (C-RM) for ID NGS Diagnostic Assays. Figure 1A illustrates a walkthrough of the proposed novel composite reference method (C-RM). Here, we show *in silico* target sequence validation with FDA-ARGOS reference genomes in combination with a wet lab validation challenge to understand the performance of ID NGS diagnostic assays. Using raw sequence data from the ID-NGS test device, *in silico* comparison of results obtained with the assay in-house database to results when using FDA-ARGOS will evaluate device bioinformatic analysis pipelines and report generation while eliminating the need for additional sample testing with a gold standard comparator (current FDA benchmarks). Overall, we anticipate the use of the C-RM method based on assay-specific subsets of clinical samples and/or microbial reference materials (MRMs) for wet lab validation and FDA-ARGOS *in silico* target sequence validation to generate scientifically valid evidence for understanding the performance of ID NGS diagnostic assays. Figure 1B lists the required quality control metrics for passing the regulatory-grade genome criteria. At a minimum, an FDA-ARGOS regulatory-grade genome adheres to six metrics (a-f). Specifically, category f details the minimum data requirements that are further described in Table 1C. In addition, Table 1D lists the 10 critical meta data that need to be ascribed to a genome to meet the regulatory-grade criteria.

Patients and clinicians need alternative solutions when conventional diagnostics (e.g., real-time PCR, culture or ELISA), fail to identify an infectious etiology. Several studies document this need of applying hypothesis-free NGS as a diagnostic of last resort, such as high-risk transplant population or failure of diagnosis with conventional diagnostics (Schlaberg, Chiu et al. 2017, Wilson, Zimmermann et al. 2017). Numerous groups have successfully applied ID-NGS technology across several unique and diverse clinical use cases. For example, isolate shotgun sequencing information uncovered unexpected transmission routes during multi-drug resistant nosocomial organism outbreaks (Snitkin, Zelazny et al. 2012, Roach, Burton et al. 2015, Snitkin, Won et al. 2017). Other studies showed use of targeted sequencing to group E. coli clonotypes from patient’s direct urine samples (Tchesnokova, Billig et al. 2013), or to detect ciprofloxacin resistance markers (Stefan, Koehler et al. 2016), resulting in antimicrobial susceptibility data and improvement in clinical outcome prediction. Finally, agnostic (unbiased, metagenomic) sequencing shows promise as a diagnostic of last resort where no other diagnostic can determine the infectious microorganism, such as the successful ID-NGS diagnosis of leptospira infection with resulting positive outcome for the patient (Wilson, Naccache et al. 2014).

ID-NGS is finding application across the infectious disease space; however, several studies document the continued need for NGS research and database curation to facilitate adoption in the clinical setting (Schlaberg, Chiu et al. 2017). Perhaps the best example, Afshinnekoo et. al. showed ID-NGS misidentification of anthrax and plague in the NYC subway system based on low quality reference genomes (Afshinnekoo, Meydan et al. 2015). A follow-up erratum by the same group (Afshinnekoo, Meydan et al. 2015) revealed the lack of evidence for biothreat organisms in these samples. This erratum attributed the anthrax misidentification to poor reference genomes leading to misattribution of toxin genes when using metagenomic data analysis tools. This lack of proper reference genomes is pervasive and represents significant knowledge gaps in public resources, thus emphasizing the necessity for targeted development of representative, accurate and well curated microbial reference genome sequences. Additional studies showed that effective use of agnostic sequencing technology, either for infectious disease identification or exclusion of infectious etiologies, is directly related to the availability of quality controlled whole-genome reference sequences (Greninger, Messacar et al. 2015, Naccache, Peggs et al. 2015, Somasekar, Lee et al. 2017). Significant efforts are still required for ID-NGS technology to transition into a routine clinical diagnostic. To facilitate this transition, prominent groups and researchers in the field have outlined steps required for proper ID-NGS use in the clinic (Gargis, Kalman et al. 2016, Schlaberg, Chiu et al. 2017, Simner, Miller et al. 2017).

In 2016, the FDA published a draft guidance for ID-NGS devices soliciting feedback on a potential regulatory pathway for targeted and pathogen-agnostic NGS diagnostic applications. This draft guidance proposed a novel regulatory strategy for ID-NGS device validation allowing wet-lab validation of an assay-specific subset of clinical samples to determine the assay preliminary diagnostic performance in combination with *in silico* validation of additional sequence targets. This *in silico* validation would entail the use of raw sequence data as an input into bioinformatic algorithms that allow a head-to-head comparison to reference genomes from the FDA dAtabase for Regulatory-Grade micrObial Sequences (FDA-ARGOS). Figure 1A illustrates this proposed novel composite reference method (C-RM). By comparison, the current regulatory paradigm relies on comparing performance of a new test device to FDA Benchmarks that are either reference standards or non-reference standards (predicate or comparison method) (See https://www.fda.gov/RegulatoryInformation/Guidances/ucm071148.htm).

To reduce some of the data generation load in clinical trials, we established and populated the FDA-ARGOS database with quality controlled microbial sequences as a tool for *in silico* target sequence validation. In this context, *in silico* target sequence validation is part of the C-RM method focused on evaluating dry lab components (bioinformatic analysis pipelines and databases) of ID NGS diagnostic assays. Using raw sequence data from the ID-NGS test device, *in silico* comparison of results obtained with the assay in-house database to results when using FDA-ARGOS will evaluate device bioinformatic analysis pipelines and report generation while eliminating the need for additional sample testing with a gold standard comparator (current FDA benchmarks). Overall, we anticipate the use of the C-RM method based on assay-specific subsets of clinical samples and/or microbial reference materials (MRMs) for wet lab validation and FDA-ARGOS *in silico* target sequence validation to generate scientifically valid evidence for understanding the performance of ID NGS diagnostic assays.

This manuscript provides our rationale and quality metrics for the FDA-ARGOS genome database initiative, outlines the need for genome gap filling in the public domain and proposes the utility of the FDA-ARGOS database resource as a novel *in silico* validation strategy for ID- NGS diagnostics.

## Materials and Methods

### FDA-ARGOS database genome deposition

Using previously identified microbe(s), nucleic acid was extracted for library preparation and sequencing. Next, microbial nucleic acids are sequenced, and *de novo* assembled using Illumina and Pac Bio sequencing platforms at the Institute for Genome Sciences at the University of Maryland (UMD-IGS). The assembled genomes were quality controlled by an ID-NGS subject matter expert working group consisting of FDA personnel and collaborators with all passing data deposited in NCBI databases. Follow this link (https://www.fda.gov/argos) for full background, collaborators and FDA-ARGOS genome status. Supplemental Table 1 lists all FDA-ARGOS genomes with accessions and statistics used in this manuscript.

### Bacterial reference genome sequencing and assembly

A hybrid sequencing approach (Koren, Schatz et al. 2012) based on long and short read NGS technology was selected using Illumina and PacBio NGS technologies to generate high quality bacterial genome sequences. Sufficient and high molecular weight genomic starting material was needed for both technologies. Sets of bacterial libraries were multiplexed on the Illumina PE HiSeq4000 using the 150bp paired-end run protocol with 24 – 48 isolates per lane. The coverage threshold was set at 300x to ensure sufficient read depth was achieved from short read NGS technology for high quality assembly generation. In addition, sets of bacterial libraries were run on the PacBio RS II P6-C4 with at least 1 SMRT cell per bacterial genome. The coverage threshold was set at 100x to ensure sufficient and economically feasible read depth was achieved from long read NGS technology for high quality assembly generation. The data were assembled both separately and in combination using a series of assembly tools, including SPAdes(Bankevich, Nurk et al. 2012), Canu (Koren, Walenz et al. 2017), HGAP (Chin, Alexander et al. 2013) and Celera Assembler (Berlin, Koren et al. 2015). Pilon (Walker, Abeel et al. 2014) was used for polishing of data. Manual curation was performed to achieve optimal assembly and consensus calling.

### Viral reference genome sequencing and assembly

Viral genome sequencing included shotgun, amplicon, and 5’/3’ RACE sequencing methods to generate full-length viral genome sequences. Sufficient and high quality genomic starting material was needed for all three approaches. Amplicon sequencing with 48 – 96 overlapping amplicons was used to generate deep coverage of known regions of the genome and was used to evaluate quasi-species in each isolate. Rapid amplification of cDNA Ends (RACE) was used to finish the 5’ and 3’ ends, and a shotgun approach generated data from all RNAs present in the sample without the level of bias present in the amplicon approach. Sets of viral libraries from all three approaches were multiplexed on the Illumina MiSeq using the 300bp paired-end run protocol. The coverage threshold was set at 100x to ensure two times amplicon coverage across the genome. The shotgun, amplicon and RACE data were assembled both separately and in combination using a series of assembly tools, including SPAdes (Bankevich, Nurk et al. 2012) and Celera Assembler (Berlin, Koren et al. 2015). Manual curation was performed to achieve optimal assembly and consensus calling.

### Calculation of FDA-ARGOS genome assembly quality control statistics

Coverage statistics were calculated for each of the FDA-ARGOS genome assemblies. Illumina coverage and PacBio coverage were calculated separately. Illumina short reads were first aligned to the assembly consensus sequence using Bowtie2 (Langdon 2015). Illumina coverage was then calculated using samtools (Li, Handsaker et al. 2009) on the resulting sam file. PacBio reads were aligned to the assembly consensus sequence using BLASR (Chaisson and Tesler 2012). PacBio coverage was then calculated using samtools (Li, Handsaker et al. 2009) on the resulting sam file. Total coverage was calculated by adding the PacBio coverage and Illumina coverage at every base pair location in the assembly consensus sequence.

### FDA-ARGOS genome annotations

Genomes were annotated with NCBI’s annotation tools to streamline the process (Angiuoli, Gussman et al. 2008, Brister, Bao et al. 2010, Klimke, O’Donovan et al. 2011, Tatusova, DiCuccio et al. 2016, Hatcher, Zhdanov et al. 2017). Bacterial sequences were annotated with NCBI’s Prokaryotic Genome Annotation Pipeline (PGAP) that combines ab initio gene prediction algorithms with homology based methods. Viral sequences were aligned with their most similar NCBI RefSeqs (NC_002549, NC_014372, NC_006432, NC_014373, NC_004162, NC_004161, NC_003899, NC_001449, NC_001544, NC_035889), using the Geneious alignment tool in the Geneious platform (Kearse, Moir et al. 2012). The setting to automatically determine detection was used, and the other parameters were set to the defaults. Gene, CDS, and mature peptide annotations from the RefSeqs were transferred to the sequences, beginning and end positions were verified for homology, and the sequences were manually reviewed for unexpected stop codons or regions of high dissimilarity. The RefSeqs used have had their annotation reviewed by NCBI curators based on available literature, and in several cases, the annotations were performed in collaboration with researchers familiar with the viruses.

### Clinical sample collection and preparation

Clinical and mock-clinical sample testing was conducted to demonstrate the utility of FDA-ARGOS. Fifteen de-identified human serum samples that were Ebola virus (EBOV) Makona positive were received from Sierra Leone; these samples were determined by the USAMRIID Office of Human Use and Ethics to be Not Human Subject Research (HP-09-32). All samples were collected and de-identified in Sierra Leone at the Kenema Government Hospital, and the samples had indirect identifiers upon receipt. Presence of virus for the human samples was determined using the previously established real-time RT-PCR assay (Trombley, Wachter et al. 2010). Samples were run in duplicate using 5µl of purified RNA on the LightCycler 480 (Roche Diagnostics Corporation). A positive sample was defined as having a quantitation cycle (Cq) value of <40 cycles with duplicate positive real-time PCR results (Table 1B).

Ten de-identified human serum samples that were suspected Bundibugyo virus positive were received from the Democratic Republic of Congo (DRC). These samples were determined by the USAMRIID Office of Human Use and Ethics to be Not Human Subject Research (HP-12-15). Presence of virus for the human samples was determined using the previously established Bundibugyo virus real-time RT-PCR assay (Trombley, Wachter et al. 2010). Samples were run in duplicate using 5µl of purified RNA on the LightCycler 480 (Roche Diagnostics Corporation). A positive sample was defined as having a quantitation cycle (Cq) value of <40 cycles (Table 1B).

One clinical *Enterococcus avium* from Children’s Hospital was used for this study and maintained at USAMRIID through the Unified Culture Collection (UCC) system. Following overnight growth of *E. avium*, (∼16 hrs), a single, isolated colony was chosen and inoculated into tryptic soy broth (ThermoFisher, Waltham MA). A glycerol stock was made from the overnight culture and colony counts were performed concurrently to determine the CFU/mL of the stock organism.

### Metagenomic and isolate shotgun sequencing

The *Enterococcus avium* sample SAMN04327393 was cultured on blood agar plates or in tryptic soy broth (ThermoFisher, Waltham MA). Samples were spiked to a final concentration of 10^5^ CFU/ml in water or whole blood matrix (BioreclamationIVT, Baltimore, MD) and 100µl was extracted using the Qiagen EZ1 viral kit (Qiagen, Valencia, CA) according to the manufacturer’s instructions. DNA concentration was quantified utilizing Qubit dsDNA BR assay kit (ThermoFisher). DNA samples were prepared for sequencing on the MiSeq platform utilizing the Nextera XT DNA library preparation kit according to the manufacturer’s instructions (Illumina, San Diego, CA). Library preparations were quantified and normalized utilizing the KAPA library quantification kit (Kapa Biosystems, Wilmington, MA) and sequenced on the MiSeq platform using the 2×150 cycle sequencing kit (Illumina). Sequencing reads were analyzed using CLC Genomic Workbench (CLC Bio, Cambridge, MA). For metagenomic analysis, paired end reads were trimmed utilizing a quality trim of 0.05 and reads below 50bp in length were removed from further analysis. Trimmed reads were then mapped to *E. avium* assembly GCF_000407245.1 and H. sapiens assembly GCA_000001405.27. Mapping parameters were as follows: mismatch costs=2, insertions costs=3, deletion costs=3, length and similarity fraction = 0.8.

### Targeted molecular inversion probe sequencing (MIPS)

The Bundibugyo virus (BDBV) and Ebola virus (EBOV) Makona clinical data samples were run using the previously described MIPS approach (Koehler, Hall et al. 2014) to capture a targeted sequence into a circular oligonucleotide. A PCR reaction and subsequent NGS on the Illumina MiSeq (2×150) amplified and identified the captured sequence using CLC genomics workbench (CLC Bio, Cambridge, MA) read mapping back to the reference genome (EBOV (GenBank # NC_002549), BDBV (GenBank # NC_014373). The percent reads classified as Bundibugyo virus or EBOV Makona was reported. The threshold for positive calls was determined by the no template control (NTC). For the MIPS approach, the remaining reads are non-specific or “junk”.

### Mock Clinical Diagnostic Evaluation

The MIPS assay was evaluated for diagnostic performance across 148 blinded samples. The limit of detection (LOD) was determined through a preliminary titration of EBOV Zaire in TRIzol starting at 10^8^ plaque forming units (pfu)/ml down to 10^2^ pfus/ml and then run in triplicate. The concentration where all three replicates yielded positive results was confirmed as the LOD across 40 replicates at that concentration. EBOV (Kikwit R4317a) in TRIzol LS was diluted to 10X (1.0E+06 pfu/ml), 5X (5.0E+05 pfu/ml) and 1X (1.0E+05 pfu/ml) LOD in triplicate in matrix also containing TRIzol LS. Nucleic acid was extracted using 400µl of each sample, along with 14 negative serum samples, on the EZ1 Virus 2.0 kit and eluted in 60µl. Presence of virus was determined with an established real-time PCR assay in triplicate for each extracted sample. Extracted RNA was amplified from 5 µl total nucleic acid using the Quantitect Whole Transcriptome Amplification Kit (Qiagen) and quantified with the Qubit dsDNA Broad Range Assay Kit. A total of 50ng cDNA was added into the MIP protocol. Library preparation was performed on the Apollo instrument using the PrepX Complete ILMN 32i DNA kit and Illumina TruSeq dual Indices. All samples were sequenced on the Illumina MiSeq using the 300 cycle kit. Sixteen samples were spiked at 10X, 5X and 1X LOD. For the mock clinical evaluation, 48 positive and 100 negative (matrix only) samples were run as described above. Threshold cutoffs for positive samples were 2X signal to noise ratio (SNR). All diagnostic performance statistics were calculated on https://www.medcalc.org/calc/diagnostic_test.php.

### Short Read Classification Using MegaBLAST Tool

The quality of the short reads was checked with FastQC. No quality trimming was conducted. We selected 100,000 short reads randomly from each of the samples (140,000 for mock clinical). The MegaBLAST function of blast+ 2.7.1 installed on FDA HPC infrastructure (https://www.ncbi.nlm.nih.gov/books/NBK153387/) was used to taxonomically classify the short reads using the default parameters Algorithm Standard Database and FDA-ARGOS database was and three databases: Algorithm Standard Database (NCBI Nt), Algorithm Standard Database and FDA- ARGOS and FDA-ARGOS alone. NCBI Nt was downloaded and constructed on 9/25/2017. The FDA-ARGOS database was constructed with FDA-ARGOS genomes (Supplemental Table 1, SAMN04327393 was excluded from the database because this reference genome was developed from the same isolate that was used as spike in material for use case 1) using the makeblastdb command. The constructed by aggregating the NCBI Nt database and FDA-ARGOS database. Default options were used to build the databases. For this study, the taxon associated with the first reported alignment was used as the taxonomic label for each read. Original MegaBLAST results were summarized to report the number of reads associated with each unique NCBI taxonomy ID called.

### Short Read Classification Using Kraken Tool

The quality of short reads was checked with FastQC. No quality trimming was conducted. We subsampled 300,000 short reads uniformly from each of the samples. Kraken 1.0 (Wood and Salzberg 2014), installed on FDA HPC infrastructure, was used to assign a taxonomic label to each short read using default parameters and three databases: Algorithm Standard Database (NCBI Nt), Algorithm Standard Database and FDA-ARGOS and FDA-ARGOS alone. NCBI Nt was downloaded and constructed on 10/5/2017. The FDA-ARGOS database was constructed with FDA-ARGOS genomes (Supplemental Table 1, SAMN04327393 was excluded from the database because this reference genome was developed from the same isolate that was used as spike in material for use case 1) using Kraken-build command. The Algorithm Standard Database and FDA-ARGOS database was constructed with both the NCBI Nt database and the FDA-ARGOS genomes. Default options were used to build the databases. For this study, the taxon associated with the first reported alignment was used as the taxonomic label for each read. Original Kraken results were summarized to report the number of reads associated with each unique NCBI taxonomy ID called.

### Short read Classification Using LMAT

The quality of the short reads was checked with FastQC. No quality trimming was conducted. LMAT version 1.2.6 (available for download at sourceforge.net/lmat, (Ames et al., 2015)), installed on Lawrence Livermore National Laboratory (LLNL) HPC infrastructure was used to assign a taxonomic label to each short read with a minimum score setting of 0.5. Match scores are calculated per read, by fitting a random null model created by simulating 1 GB of random sequence for each model dependent on read length and GC content. Three databases, the Algorithm Standard Database (LMAT DB), the stand-alone FDA-ARGOS database (Supplemental Table 1, SAMN04327393 was excluded from the database because this reference genome was developed from the same isolate that was used as spike in material for use case 1) and an aggregated database consisting of both the LMAT DB database and the stand-alone FDA-ARGOS database were used. LMAT results were summarized to report the number of reads associated with each unique NCBI taxonomy ID.

## Results

### Filling gaps in public resources with targeted reference genomes

In 2013, FDA in collaboration with the Department of Defense (DoD) and the National Center for Biotechnology Information (NCBI) assessed the quality and diversity of sequenced microbial genomes present in public databases. A majority of pathogens appeared to be represented by multiple entries, however, many of these genomes were incomplete or of unknown quality. In fact, a thorough examination of the entire public domain revealed some pathogens were underrepresented or completely absent. Our 2013 review, supported by several publications (Schatz and Langmead 2013, Land, Hyatt et al. 2014, Land, Hauser et al. 2015), revealed biased phylogenetic coverage usually attributable to research funding for specific microbial model organisms. At the time, NCBI GenBank covered less than 8,000 bacterial and archaeal genome sequences with at least half submitted by the four largest genome sequencing centers: Broad Institute, DOE Joint Genome Institute, Institute for Genome Sciences and TIGR/JCVI. Additionally, many sequences lacked accompanying metadata and raw read information. These issues provided the impetus for *de novo* generation of FDA-sponsored reference sequences of the highest quality achievable using state-of-the-art genomic sequencing technologies (Koren, Schatz et al. 2012). With this effort, FDA intended to establish quality control metrics for microbial genomes that could be used in ID-NGS test validation. Only genomes with the highest technically achievable quality would qualify as regulatory-grade genomes. Factors essential to reach that goal were: 1) knowledge of the technology used to generate the sequences, 2) access to raw sequence information to reproduce the data, and, 3) access to relevant metadata. Perhaps the most significant missing piece of information for previously generated reference genomes was the lack of an independent reference method that reliably linked the microbial organism identification to the sequence data. In this context, qualification of microbial reference genomes requires organism identification with a recognized reference method as this remains a primary requirement for validation of a new diagnostic device.

FDA, DOD, NCBI and other agencies using scientific literature, a phylogenetic data mining approach, and FDA microbial species-specific guidance documents identified more than 1000 gaps in public microbial genomic repositories. We prioritized these gaps and selected biothreat microorganisms, common clinical pathogens and closely related species (See Supplemental Materials for the organism gap list). The primary objective of this regulatory science research and tool development effort centered on the generation of an initial set of 2000 quality-controlled microbial FDA-ARGOS reference genomes. These genomes are generated with a hybrid assembly approach using short and long read sequencing technologies (Koren, Schatz et al. 2012). An initial collection criterion focused on sequencing at least 5 diverse isolates per species to cover temporal and spatial genome plasticity and initiate the construction of a regulatory-grade microbial genome model.

### FDA-ARGOS, what’s that?

FDA and collaborators established the publicly available database, FDA dAtabase for Regulatory-Grade micrObial Sequences (FDA-ARGOS), to fill these defined gaps for genomic sequences. Here, we present the first subset of 487 FDA-ARGOS genomes with NCBI accessions (Figure 2, Supplemental Table 1). Of the 487 isolates, 88.3 percent were bacteria, 11.1 percent were viruses and 0.6% were eukaryotes, representing 189 different taxa. In total, 81.9 percent of genomes were of clinical origin with the remaining 18.1 percent environmental genomes from closely related species near-neighbors (Supplemental Table 2). Over 500 isolates are currently being sequenced and at different stages in the FDA-ARGOS genome generation pipeline.

**Figure 2:**
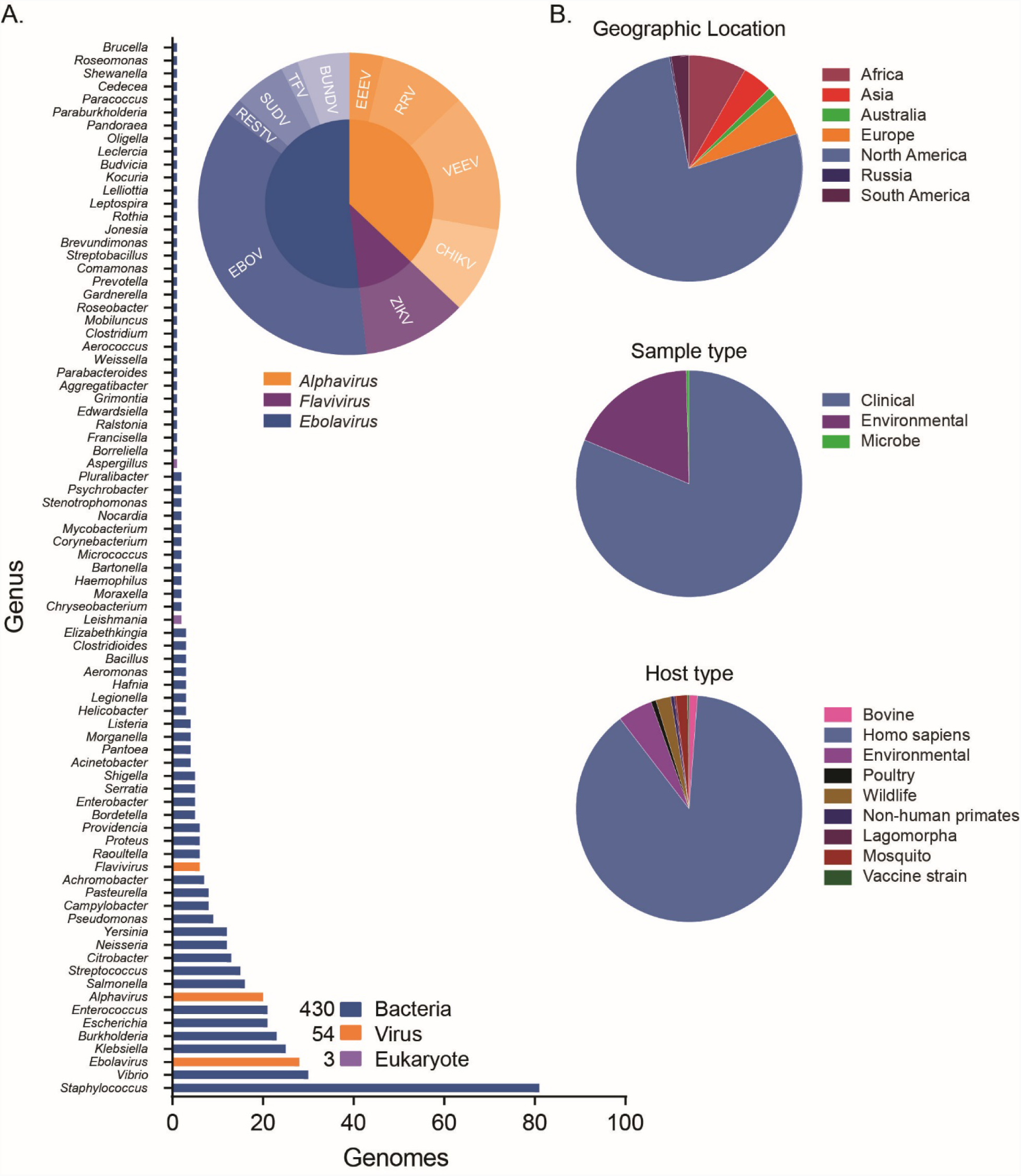
FDA-ARGOS Reference Genome Database. Summary statistics of the current 487 microbial genomes show primary coverage of FDA- ARGOS resides with bacterial isolates, followed by viruses and then eukaryotic parasites (A). Supplemental Table 1 provides accessions for all 487 genomes currently available publicly. A majority of FDA-ARGOS constituents (B) originate from North America and are from human clinical isolation.

Use of advanced sequencing technologies (Koren, Schatz et al. 2012) helped define the characteristics for regulatory-grade genomes. Specifically, Figure 1B provides a summary of required FDA-ARGOS metrics to support a determination of regulatory-grade genome. All FDA- ARGOS genomic submissions demonstrated: 1) organism identification prior to sequencing by a recognized independent reference method, 2) sequence generation with at least two sequencing methodologies (e.g., long read and short read NGS), and, 3) *de novo* assembly with high-depth of base coverage. Each microbial isolate assembled genome sequence conformed to a minimum of 95 percent coverage with 20X depth at every position while also providing concordant NCBI taxonomy-specific average nucleotide identity (ANI) thresholds for microbial organism identification (Ciufo, Kannan et al. 2018) with independent identification methods. All FDA-ARGOS samples were concordant between *de novo* sequencing identification and independent organism identification method (Supplemental Table 2 lists independent identification method data).

As mentioned above, hybrid error-correction with long and short read sequencing technology was considered for establishing minimum FDA-ARGOS regulatory grade data requirements. Figure 1C outlined these criteria and included sample name, 10 meta data fields (based on NCBI BioSample submission requirements), raw reads, assemblies with coverage, N50, L50 and annotations. Importantly, FDA-ARGOS genomes are tied to a minimum of 10 critical sample metadata fields (Figure 1D): independent organism confirmation by recognized reference method, culture collection, and, the following required NCBI BioSample fields: organism, strain, isolation source, host, collected by, taxonomy ID, contact and package information. Supplemental Table 2 shows metadata coverage metrics for all 487 FDA-ARGOS genomes. The 10 sample metadata fields are 100 percent completed and available throughout the sample set with 5 additional metadata metrics are recommended, such as geographic location, collection date, host disease, host sex and host age (BioSample documentation https://www.ncbi.nlm.nih.gov/biosample/docs/attributes/). In terms of clinical representation, 81.9 percent of clinical samples in the collection are associated with known phenotype/host disease.

Critical for the designation of genomes as ‘regulatory-grade genomes’, was the institution of quality control metrics for all aspects of the genome generation. To objectively identify such quality control metrics, we performed internal quality control assessments of all 487 genome assemblies (See methods for calculation of FDA-ARGOS genome assembly quality control statistics, Supplemental Table 1). Figure 3 shows the quality of FDA-ARGOS genome assemblies compared to the representative 2013 NCBI GenBank database and the representative 2018 NCBI GenBank database. Both, the 2013 and 2018 NCBI database captures held up to 50 NCBI assemblies for each species within the FDA-ARGOS database from the respective year. In relative number of assemblies, 2018 NCBI database contained 3535 while the 2013 contained 1617. Overall, we observed higher quality in the FDA-ARGOS genome dataset for the coverage, N50, and L50 quality assembly metrics compared to the 2013 and 2018 NCBI GenBank public genome dataset (Figure 3 A, B and C respectively). Figure 3D demonstrated that only 675 out of the 3535 2018 NCBI GenBank assembled genomes, or 20 percent, showed comparative assembly quality to FDA-ARGOS genome sequences when considering one of the reported assembly quality metrics. More importantly, when considering all quality control assembly metrics, only 11 out of the 3535 2018 NCBI GenBank assembled genomes, or 0.3 percent, showed comparable quality to FDA-ARGOS genome assemblies.

**Figure 3:**
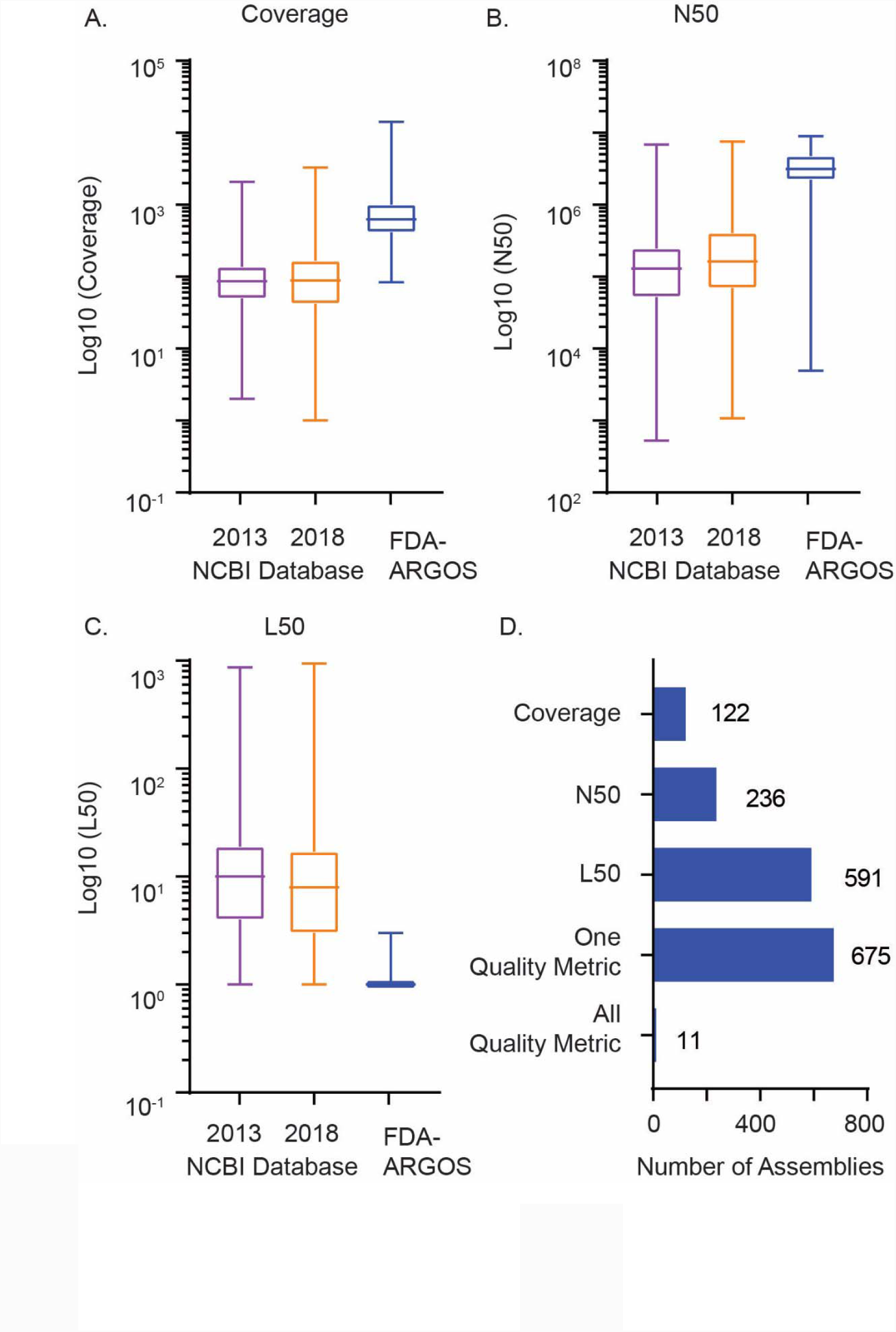
FDA-ARGOS Reference Genome Assemblies Quality Metrics. Comparative microbial genome assembly quality metrics contrasted current FDA-ARGOS assemblies to 2013 and 2018 NCBI GenBank assemblies submitted for each species captured within the FDA-ARGOS database. Assembly quality metrics measured included: (A) median coverage, (B) median N50, (C) median L50 and (D) number of 2018 NCBI genomes that exhibited all, one or a specific quality control metric used to vet FDA-ARGOS genomes for inclusion. The NCBI assemblies were downloaded on August 6, 2018.

We expect refinement of the quality metrics for ‘regulatory-grade’ genome status (Figure 1B) as we continue to populate the FDA-ARGOS with additional quality-controlled genomes; therefore, we established the requisite that all genomes should be available publicly. Deposition for all FDA-ARGOS genomes requires that raw reads, assembled genomes and associative metadata are publicly available (https://www.ncbi.nlm.nih.gov/bioproject/231221). (Check https://fda.gov/argos for additional background information and updated genomes).

### FDA-ARGOS fills critical gaps in public sequence repositories - Use Case 1: Enterococcus avium

Several regulatory science research considerations arose during the process of generating FDA-ARGOS genomes, including the initial impetus for this effort, gap filling. Our first use case documented the importance of genome gap filling with FDA-ARGOS quality-controlled genomes, and the impact of lack of publicly available genomes for medically important microbes on potential diagnostic applications. Specifically, we tested whether the addition of quality-controlled reference sequences into the public repositories impacted the NGS pathogen detection of a metagenomic shotgun sequencing approach of a mock clinical *E. avium*-spiked human blood sample at clinically relevant titers. An isolate from reference genome SAMN04327393, which was removed from reference databases for data analysis, was used as a mock clinical E.avium sample. Initial read mapping using CLC Genomics and *E. avium* sequences from publicly available databases as a reference demonstrated *de novo* assembly of *E. avium* data was not possible due to only an average 424.4 mapped paired end reads (Supplemental Table 7). For frame-of-reference, we would need over 30,000 reads to *de novo* assemble an entire genome of approximately 5 Mb at 1X coverage, assuming a read size of 150 bp and perfect quality of each generated read at all positions.

Subsequent bioinformatics data analysis of the *E. avium* metagenomics shotgun paired-end reads data showed the critical gap filling and utility of the FDA-ARGOS database resource. We analyzed the effect of genome gap filling with MegaBLAST (Morgulis, Coulouris et al. 2008) and Kraken (Wood and Salzberg 2014) by determining the number of *E. avium* reads classified from the mock clinical human blood sample with and without FDA-ARGOS genomes used in the respective bioinformatics tools reference databases. Intuitively, a majority, over 98 percent, of approximately 12 million paired-end reads for each replicate sample mapped against the human genome with only 2 percent or less mapping to non-human sequences with both algorithms (Figure 4A). In contrast, application of MegaBLAST and Kraken with FDA-ARGOS alone yielded zero human reads due to the lack of human reference in that database. Reads classified as *E. avium* ranged from an average 3829 and 840 when FDA-ARGOS genomes were added to the algorithm reference database compared to an average 29 and 0 reads when these genomes were absent for MegaBLAST and Kraken, respectively (Figure 4B, Supplemental Table 4). Interestingly, while *E. avium* genomes were available in the NCBI Nt database and part of the read classification for MegaBLAST analyses, positive ID-NGS identification required the addition of quality-controlled FDA-ARGOS reference genomes. MegaBLAST tool with FDA- ARGOS data as the standalone reference database generated the largest effect with an additional 1495 *E. avium* reads classified. MegaBLAST classified additional reads with the stand- alone FDA-ARGOS database most likely because the quality-controlled *E. avium* genomes were not mixed with lower quality genomes in the standard algorithm database as competition with lower quality and closely related genomes removed.

**Figure 4.**
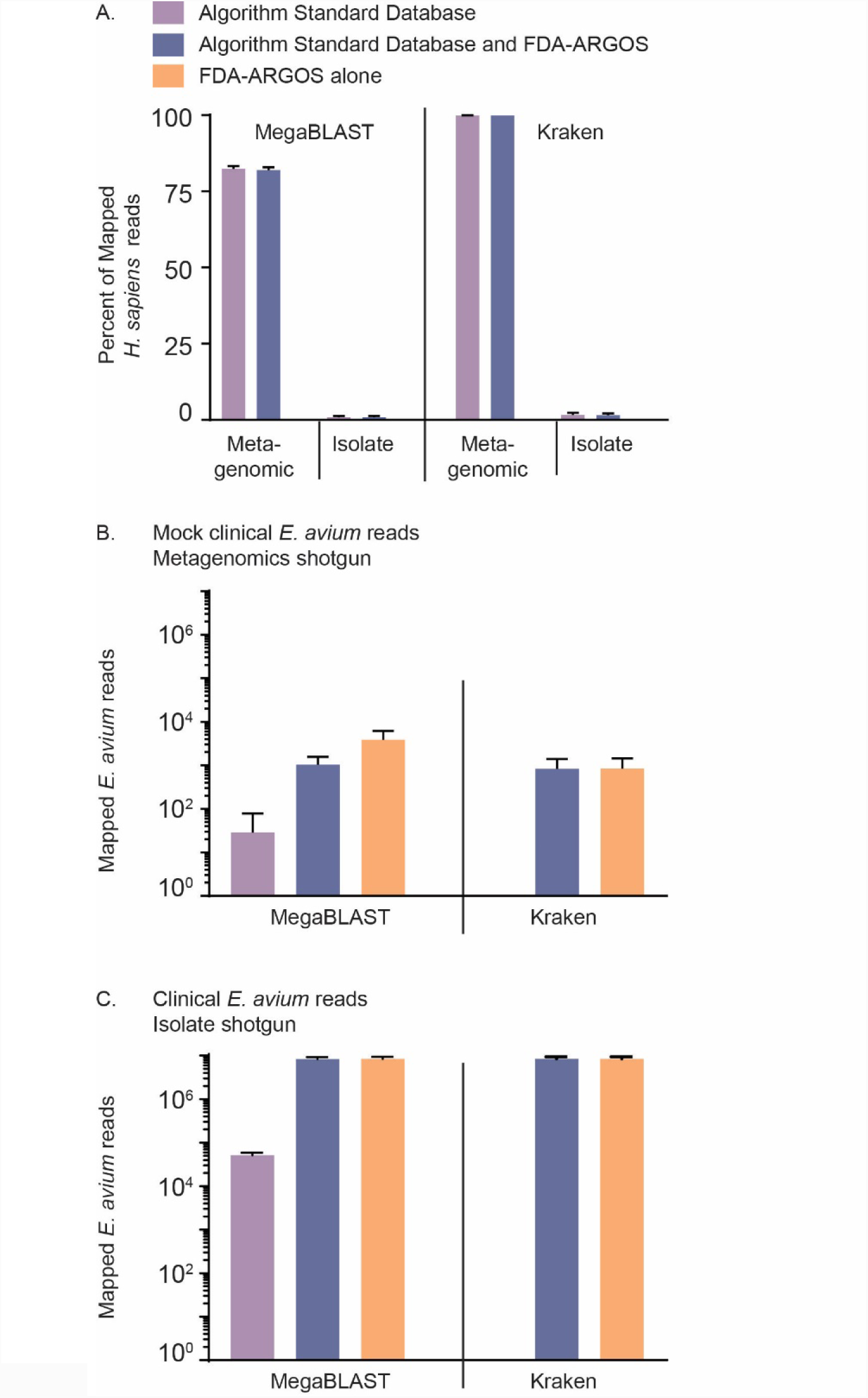
Read Classification Results from Shotgun Sequencing for Identification of *Enterococcus avium*. Visualizing sample analyzed with both MegaBLAST and Kraken percent human reads mapped (A)from metagenomics and isolate sequencing shotgun sequencing showed sequencing clinical matrix resulted in a majority of reads mapping to host background. The number of *E. avium* reads correctly classified from metagenomic samples using three different reference databases(B)varied based on whether MegaBLAST and Kraken standard databases, standard database plus FDA-ARGOS, or FDA-ARGOS alone was used for read mapping. Evaluation of *E. avium* reads for just the clinical isolate without matrix (C) resulted in a similar relationship of greater number of reads mapped when FDA-ARGOS genomes were used in comparison to the algorithm standard reference database. All Reference Data Sets and the FDA-ARGOS Database are publicly available for data analysis and tool comparison.

Finally, we performed *E. avium* isolate shotgun sequencing without clinical matrix to obtain sufficient data to illustrate the critical nature of having quality-controlled reference genomes. Using the aforementioned bioinformatics tools, data analysis showed the impact of read classification solely focused on *E. avium* and determined if the addition of FDA-ARGOS genomes to public databases affected read mapping. Figure 4C shows that addition of FDA- ARGOS *E. avium* reference genomes significantly increased read classification performance based on the number and percent of *E. avium* reads classified (Figure 4C, Supplemental Table 5). On average, for E.avium isolate shotgun sequencing, 8,406,630 reads out of a total 12 million reads classified as *E. avium* when the FDA-ARGOS database resource upon addition to the algorithm standard reference database (NCBI Nt) compared to 25,800 reads without FDA- ARGOS added. Amalgamation of FDA-ARGOS genomes into standard sequence reference databases resulted in *E. avium* contributing between 84 to 96 percent of the total reads classified (Figure 4C). Interestingly, top hits from the MegaBLAST tool using NCBI Nt database (containing 4 *E. avium* genomes but not at regulatory grade quality, Supplemental Table 3) showed over 10 percent of total classified reads mapped to ‘Bos Taurus’ or ‘Enterococcus faecium’. These top hits were potentially database contaminants and illustrate the risk of using non-curated databases in ID-NGS diagnostics. Data analysis with the Kraken tool and the algorithm standard reference database (NCBI Nt) resulted in 0 mapped reads because the Kraken tool reference database lacked *E. avium* genomes (Figure 4C).

For future benchmarking efforts of bioinformatics tools, we provide all *E. avium* Data Sets (Supplemental Material).

### In Silico Comparison: Regulatory-grade genomes are sufficient for Ebolavirus Target Sequence Validation – FDA-ARGOS Use Case 2

A major incentive for the development of FDA-ARGOS was to enable and promote innovation for ID-NGS medical devices. Through the process of populating the FDA-ARGOS database, the concept of partial *in silico* validation, rather than completely empirical validation of clinical trial samples with an independent gold standard reference method, matured. We chose FDA-ARGOS Ebola reference sequences (Supplemental Table 1) and a targeted ID-NGS assay, the Ebola virus molecular inversion probes (MIPS), to evaluate the application of FDA- ARGOS as an *in silico* target sequence validation tool. Table 1 showed the diagnostic performance of the MIPS ID-NGS assay with clinical Bundibugyo virus and EBOV Makona samples reported as a more sensitive assay, EBOV Real-Time PCR (RT-PCR) assay (Trombley, Wachter et al. 2010). When assessing 10 clinical Bundibugyo virus and 15 clinical EBOV Makona samples, concordant real-time PCR and MIPS positive results ranged from 9 out of 10 clinical samples (Table 1A) to 6 out of 15 (Table 1B), respectively. Intuitively, lower quantitation cycle (C_q_) values correlated with higher MIPS read classification, suggesting the capability of ID-NGS to detect organisms was dependent on the starting concentration of the target genomic material. MIPS false negative calls for low target analytes suggested that complete *in silico* validation is an unrealistic approach for clinical trials without comparison to some gold standard reference method, in this case real-time PCR.

**Table 1A:** Diagnostic Benchmark and *In Silico* Target Sequence Validation with FDA-ARGOS: Bundibugyo Performance Summary.

| Sample | Real-Time PCR (Benchmark) | MIPS (Test Device) | FDA-ARGOS (*In Silico* Target Sequence Validation) |  |
| --- | --- | --- | --- | --- |
|  | Quantitation Cycle (Cq) Value | Percent Classified | MegaBLAST Percent Classified | Kraken Percent Classified |
| 2012-1 | 22.97/22.95 | 54.95% | 59.84% | 70.89% |
| 2012-16 | ND | 0.02% | 0.76% | 0.02% |
| 2012-91 | ND | 0.03% | 0.65% | 0.04% |
| 2012-95 | ND | 0.02% | 0.59% | 0.03% |
| 2012-99 | ND | 0.02% | 0.67% | 0.06% |
| 2012-120 | 23.46/23.38 | 41.87% | 45.14% | 57.56% |
| 2012-147 | 25.58/25.52 | 27.23% | 29.78% | 47.52% |
| 2012-153 | 28.14/27.96 | 38.30% | 40.97% | 50.70% |
| 2012-176 | 37.01/36.54 | 0.01% | 0.87% | 0.01% |
| 2012-198 | ND | 0.02% | 0.71% | 0.03% |
| NTC | N/A | 0.02% | 1.59% | 1.05% |

Illustration of an application of the *in silico* target sequence validation method to a targeted ID sequencing assay (MIPS). Two bioinformatics tools (MegaBLAST and Kraken) were selected to classify reads using default parameters utilizing FDA-ARGOS as a standalone reference database. Table 1A showed the traditional benchmark comparison of the MIPS assay to Real-Time PCR (RT-PCR) results. Benchmark positive values were only noted for samples that yielded duplicative positive results by RT-PCR. Percent reads classified only refer to percentage of reads that were assigned to Bundibugyo or Ebola virus, the remaining reads are non-specific or "junk".

**Table 1B:** Diagnostic Benchmark and *In Silico* Target Sequence Validation with FDA-ARGOS: Ebola Makona Performance Summary.

| Sample | Real-Time PCR (Benchmark) | MIPS (Test Device) | FDA-ARGOS (*In Silico* Target Sequence Validation) |  |  |
| --- | --- | --- | --- | --- | --- |
|  | Quantitation Cycle (Cq) Value | Percent Classified | MegaBLAST Percent Classified | Kraken Percent Classified | LMAT Percent Classified |
| 3754-2 | 35.11/35.72 | 0.05% | 0.60% | 0.06% | 0.03% |
| 3754-4 | 33.83/33.36 | 0.06% | 0.68% | 0.06% | 0.03% |
| 3811-2 | 36.17/36.14 | 0.07% | 0.56% | 0.08% | 0.04% |
| 3856-1P | 15.95/15.98 | 76.63% | 80.91% | 79.50% | 42.48% |
| 3913-5 | 34.00/33.77 | 0.00% | 0.68% | 0.00% | 0.01% |
| 3958-4 | 32.77/33.28 | 0.04% | 0.65% | 0.05% | 0.05% |
| 3991-2 | 33.92/33.62 | 0.00% | 0.65% | 0.00% | 0.01% |
| 4007-2 | 26.30/26.55 | 21.33% | 22.41% | 21.81% | 12.12% |
| 4015-1 | 21.66/21.66 | 74.87% | 76.35% | 75.82% | 39.35% |
| 4033-1 | 16.59/16.32 | 76.64% | 79.59% | 78.97% | 41.79% |
| 4268-1P | 25.05/25.15 | 29.71% | 30.43% | 29.97% | 15.87% |
| 4468-3 | 35.78/35.91 | 0.04% | 0.59% | 0.05% | 0.03% |
| 4641-3P | 31.81/31.82 | 0.03% | 0.59% | 0.04% | 0.02% |
| 4726-1 | 21.22/21.21 | 53.95% | 56.44% | 54.28% | 30.05% |
| 4845-3 | 35.17/36.71 | 0.00% | 0.66% | 0.01% | 0.01% |
| NTC | N/A | 0.02% | 0.86% | 0.04% | 0.01% |

Illustration of an application of the *in silico* target sequence validation method to a targeted ID sequencing assay (MIPS). Three bioinformatics tools (MegaBLAST, Kraken and LMAT) were selected to classify reads using default parameters utilizing FDA-ARGOS as a standalone reference database. Table 1B showed the traditional benchmark comparison of the MIPS assay to Real-Time PCR (RT-PCR) results. Benchmark positive values were only noted for samples that yielded duplicative positive results by RT-PCR. Percent reads classified only refer to percentage of reads that were assigned to Bundibugyo or Ebola virus, the remaining reads are non-specific or "junk".

Consistent concordance between the benchmark RT-PCR assay, the MIPS test device and the FDA-ARGOS *in silico* target sequence validation was important for establishing confidence in considering *in silico* comparison method for clinical sample ID calling. To test this assumption, we used three bioinformatics tools, MegaBLAST (Morgulis, Coulouris et al. 2008), Kraken (Wood and Salzberg 2014) and LMAT (Ames, Hysom et al. 2013) to evaluate the proposed *in silico* target sequence validation method (Figure 1A), and to verify the potential for using *in silico* comparison without any empirical validation. MegaBLAST and Kraken analyses of raw sequence data for Bundibugyo virus samples using the three different read classification tools in combination with FDA-ARGOS as the reference genome database showed complete agreement for MIPs and *in silico* calls (Table 1A). Because the *in silico* comparison missed the classification call against the gold standard PCR benchmark test for a sample with low analyte levels (1 false negative result for the *in silico* validation), we performed a more in depth analysis of the additional EBOV Makona samples across three bioinformatics tools, MegaBLAST, Kraken and LMAT (Table 1B). These analyses showed similar results to the Bundibugyo virus data at 100% agreement with test device, but only for samples with low C_q_ or high input concentrations of the target organism. Additional analyses comparing results for each bioinformatics tool reference databases with and without FDA-ARGOS genomes added, produced similar results demonstrating that FDA-ARGOS alone was sufficient for *in silico* comparison (Supplemental Table 6). Overall, these data suggested *in silico* sequence comparison would be completely reliant on the inherent sensitivity of the sequencing assay to generate sequence read data for comparison, therefore Composite Reference Method (C-RM) (combining *in silico* sequence comparison with a wet lab validation challenge) is necessary for full validation of the test ID- NGS device. Figure 1A illustrates the proposed novel C-RM, highlighting this need for empiric assessment of an ID-NGS assay-specific subset of samples or well defined microbial reference materials.

Evaluation of the clinical samples suggested a need for benchmarking ID-NGS assays to currently implemented reference methods, thus the application of the C-RM. To document the application of MIPS Ebola Makona ID-NGS assay benchmarking, we performed a mock clinical trial to assess the assay-specific wet-lab subset evaluation as part of the proposed C-RM. Initially, we performed a preliminary limit of detection (LOD) evaluation to determine the scope of the mock clinical evaluation. These experiments showed a preliminary LOD of 10^5^ with linear dose response correlation to EBOV input across the titration (Supplemental Table 8). An additional 40 positive replicates performed on two independent days, two independent runs confirmed the LOD at 10^5^ pfu/ml for EBOV. This concentration formed the basis for spike-in levels of the mock clinical trial. From a total of 148 samples tested, 48 constituted positive spiked samples with 16 at high (10x LOD), 16 at medium (5x LOD) and 16 at 1x LOD for the MIPS assay (Table 2). In this mock clinical trial, all spiked samples were positive via real-time PCR (data not shown). Only 9 out of 16 samples at 1X LoD for the MIPS assay were positive with 37 out of 48 samples positive across the entire sample set in this analysis. However, the positive predictive value (PPV) and negative predictive value (NPV) for the MIPs assay were: 97.4% and 90%, respectively at or above the limit of detection with a prevalence of 32.4%. In addition, Table 2 lists the positive and negative predictive values for prior probabilities of infection from 0-1. The PPV and NPV metrics are important predictive analytics tools to provide performance characteristics for how the ID-NGS diagnostic test will perform in a clinical context. These data provide a rationale for developers using partial *in silico* validation when false negative rate is low.

**Table 2:** Mock Clinical Evaluation of EBOV NGS Performance.

A. Experimental design and results
| PFU/ml | n | Avg EBOV Reads | Avg %Reads Mapped | CoV | Positive Samples | Negative Samples |
| --- | --- | --- | --- | --- | --- | --- |
| 1000000 (10X) | 16 | 5442.5 | 2.66% | 136.55% | 15 | 1 |
| 500000 (5X) | 16 | 2777.5 | 2.49% | 152.33% | 13 | 3 |
| 100000 (1X) | 16 | 351.5 | 0.58% | 247.57% | 9 | 7 |
| NTC | 100 | 4 | 0.00% | 571.69% | 1 | 99 |

B. Diagnostic performance statistics
| N | Positive Predictive Value | Negative Predictive Value | Sensitivity | Specificity | Prevalence |
| --- | --- | --- | --- | --- | --- |
| 148 | 97.37%<br>(83.95% to 99.62%) | 90.00 %<br>(84.26% to 93.80%) | 77.08%<br>(62.69% to 87.97%) | 99.00%<br>(94.55% to 99.97%) | 32.43%<br>(24.98% to 40.61%) |

C. Diagnostic performance statistics for prior probabilities
| Prior probability of infection | Positive Predictive Value | Negative Predictive Value |
| --- | --- | --- |
| 0 | 0 | 1 |
| 0.01 | 0.44 | 1 |
| 0.05 | 0.8 | 0.99 |
| 0.1 | 0.9 | 0.97 |
| 0.15 | 0.93 | 0.96 |
| 0.2 | 0.95 | 0.95 |
| 0.25 | 0.96 | 0.93 |
| 0.3 | 0.97 | 0.91 |
| 0.4 | 0.98 | 0.87 |
| 0.5 | 0.99 | 0.81 |
| 0.6 | 0.99 | 0.74 |
| 0.7 | 0.99 | 0.65 |
| 0.75 | 1 | 0.59 |
| 0.8 | 1 | 0.52 |
| 0.85 | 1 | 0.43 |
| 0.9 | 1 | 0.32 |
| 0.95 | 1 | 0.18 |
| 0.99 | 1 | 0.04 |
| 1 | 1 | 0 |

Demonstration (A) of the preliminary diagnostic performance (B) of a targeted ID sequencing assay (MIPS) during a mock clinical trial using 48 positive Ebola samples and 100 Ebola negative samples. Numbers in parentheses represent the 95% Confidence Interval. Positive and negative predictive values are shown for prior probabilities (C) of infection ranging from 0-1.

For future benchmarking efforts of bioinformatics tools, we have provided all Ebola Data Sets (Supplemental Material).

## Discussion

To encourage innovation and support the infectious disease community, we provide here the FDA-ARGOS resource as a tool for ID-NGS assay development, reference database and *in silico* target sequence validation as part of a novel Composite Reference Method (C-RM). This manuscript describes the database, specifically highlighting: 1) the quality metrics of regulatory-grade genomes for database inclusion, 2) benefits of FDA-ARGOS in filling pathogen genome knowledge gaps for device output, and 3) describes some use cases for FDA-ARGOS.

A critical aspect for assessing performance of any diagnostic is the availability of minimum quality control metrics for data, genomic or otherwise, for validation. Defined here are the FDA- ARGOS ‘regulatory-grade’ genome criteria that provide ID-NGS diagnostic assay developers and the scientific community with traceable and quality-controlled genomes. These high-quality genomes coupled with a streamlined approach for comprehensive expansion of FDA-ARGOS beyond the initial 2000 genomes is essential for continued ID-NGS diagnostic assay development.

FDA-ARGOS genome sequencing and research resulted in six broad quality metrics (Figure 1B) defining ‘regulatory-grade’ genome criteria required for current and future FDA-ARGOS contributors. All extant genomes in the FDA-ARGOS database (Supplemental Table 1) adhere to the quality metrics of 95% coverage with 20X depth at every position across the entire assembled genome. This metric applies to the initial deposition of, minimally, 5 genomes of any genus/ species added to FDA-ARGOS. These 5 genomes define the “FDA-ARGOS core genome”. After 5 or more regulatory grade genomes per genus/species are available in the database, we will consider lower threshold metrics for FDA-ARGOS inclusion to capture novel and/or unique genomes that may be diagnostically informative. Future efforts will apply these metrics to existing genomic information in the public domain coupled with deep learning methods and artificial intelligence to inform an external genome qualification tool greatly expanding utility of the FDA-ARGOS database.

Lack of high quality reference genomes challenges the accuracy of ID-NGS identification for queryable microbial pathogen. The genome gap filling use case with regulatory-grade *E. avium* genomes highlights current challenges with infectious disease NGS technology when using minimal-, non-curated or absent reference databases. The end result potentially leading to the lack of a diagnostic call or even misdiagnosis. These data were punctuated by two key findings: 1) *de novo* assembly of the data was not possible due to the low number of reads in clinical matrix and 2) limited *E. avium* species reference genomes in publicly available databases made the sample identification almost impossible (Supplemental Table 3). The latter point is extremely relevant for the intent of ID-NGS for diagnostic applications. In the case presented here, the top microbial sequence hit did not equate to the microbe of interest due to lack of representation in the reference database. Intuitively, addition of FDA-ARGOS and relevant genomes mitigated this issue. In addition, *E. avium* isolate sequencing results showed the dependency of both classification method (such as MegaBlast and Kraken) and database used. This last aspect of the *E. avium* use case informed the consideration of the C-RM and opened the possibility for utilizing a suite of validated bioinformatics tools for *in silico* target sequence validation.

There are two basic contrasting philosophies in circulation regarding genomic information and ID-NGS: 1) all information, whatever the quality, is useful towards making a diagnosis, the more data the better, with the assumption of diagnosis relying on error correction through iteration, or, 2) quality-controlled, highly curated genomes are required as a solid foundation, more information is better, however, diagnostics require quality-controlled genomes to inform the basis of diagnosis. Experiments and data presented here support the latter of these two arguments. Specifically, while *E. avium* reference genomes were available in NCBI Nt database and were part of the read classification for MegaBLAST analyses, positive ID-NGS identification of *E. avium* required the addition of quality-controlled FDA-ARGOS reference genomes. In addition, read mapping of isolate shotgun data, without any clinical matrix, showed indeterminate results for *E. avium* without FDA-ARGOS in contrast to 80 percent of total reads mapped as *E. avium* upon addition of these regulatory-grade genomes to the reference database. A similar increase in performance in *E. avium* reads classified resulted when using FDA-ARGOS *E. avium* reference genomes for metagenomics shotgun data, in whole blood, even with human reads occupying >98% of sequencing real estate.

Quality and coverage of targeted organisms are critical aspects for ID-NGS transition into the clinical space; however, to foster the transition, new methods are required to lessen the burden for validating ID-NGS against all queryable pathogens. This manuscript documents methods for use of FDA-ARGOS reference genomes in *in silico* sequence comparison as part of the proposed novel C-RM. We showed here that the *in silico* validation of Bundibugyo virus and Zaire ebolavirus can use FDA-ARGOS genomes as the comparator. For MIPS positive samples, there was 100 percent concordance between the gold standard real-time PCR comparator, and the *in silico* comparison. This supports the feasibility of implementing this strategy to shorten future clinical NGS-based assay evaluation studies. A potential mitigation for this issue, where real-time PCR was more sensitive than the MIPS NGS assay especially at high C_q_ values, is the application of additional enrichment strategies to bring ID-NGS to similar sensitivities as the gold standard (Briese, Kapoor et al. 2015, O’Flaherty, Li et al. 2018). However, in the current form, observed lower sensitivity of the MIPS assay compared to real-time PCR shows the necessity for a C-RM and incorporating additional empirical studies, i.e., an assay-specific subset of clinical samples going through wet-lab comparison as part of the clinical validation. Discordant results at high C_q_ values highlight the perils of solely applying *in silico* sequence comparison. Without any empirical evaluation, *in silico* comparison would only provide results within the sensitivity ranges of the test ID-NGS device without providing the needed benchmark for sensitivity compared to a gold-standard such as real-time PCR. Therefore, as part of the C- RM, we demonstrate a preliminary performance assessment a against a gold-standard for a subset of the clinical trial samples with the intent that the remainder of the clinical trial samples could be validated via *in silico* sequence comparison. Different sample read depths may be required to achieve the desired identification performance for various organisms. Assay developers might be required to use an external comparator only for *in silico* validation results where the test device and *in silico* comparison yielded a discordant result. We envision this C- RM to be a primary utility of the FDA-ARGOS genome database tool for medical device development. We hope that FDA-ARGOS will spur innovation and expedite regulatory science, and ultimately enable ID-NGS as a diagnostic to enter the clinic.

The FDA-ARGOS reference genome resource is a constantly evolving public database instance and intended to mature over time with community support and genomic technology advancements. Continued population and expansion of the FDA-ARGOS database resource will be required to cover the panoply of infectious microorganisms. In this proposed *in silico* validation with FDA-ARGOS, the need for comprehensive regulatory-grade genome coverage is clear, however, no one entity can perform all the needed sequencing. We are therefore working on a pathway for external genome qualification to streamline and expand FDA-ARGOS resource as needed. Both the external genome qualification and continued research to apply this regulatory-grade standard to unculturable and emerging pathogens will be the focus of future research.

Further population and curation of the database will support the success of FDA-ARGOS and promote adoption by the NGS community. The FDA-ARGOS team openly invites additional collaborators from the scientific community to assist in filling the gaps in this public resource. FDA-ARGOS and collaborators are specifically searching for unique, hard to source microbes such as biothreat organisms, emerging pathogens, and clinically significant bacterial, viral, fungal, and parasitic genomes. As stated, the goal is to collect sequence information for a minimum of 5 isolates per species and we solicit any potential collaborators interested in supplying these 5 isolates for gap-filling to contact and authors of this paper. For more information about contributing samples for UMD-IGS sequencing as part of FDA-ARGOS efforts, or to qualify existing genomes by the FDA, please.

## Supporting information

## Acknowledgements

- This project has been funded with Federal funds from the Office of Counterterrorism and Emerging Threats, Food and Drug Administration, Department of Health and Human Services, under Contract No. HHSF223201310109C, HHSF223201510106C, HHSF223201610073C and the Department of Defense, under Contract No. 224-15- 6506R.
- This research was supported by the Intramural Research Program of the NIH, National Library of Medicine.
- This project was funded by DTRA, Contract No. CB10245.
- Sample contributions for the initial set of 500 from the U.S. Army Medical Research Institute of Infectious Diseases, the Department of Defense Critical Reagents Program, Public Health Agency Canada, Public Health England, the University of Texas Medical Branch, BC Centre for Disease Control, American Type Culture Collection, Rockefeller University, FDA-CBER (Maria Rios, Robert Duncan, Rafaelle Gusmao), FDA-CFSAN (Eric Brown, Marc Allard, Maria Hoffman, Cary Pirone), FDA-CVM (Patrick McDermott, Shaohua Zhao), Children’s National Hospital (Joseph Campos, Brittany Goldberg, Chelsie Geyer), University of Colorado School of Medicine (Thomas Morrison, Sudhakar Agnihothram)
- The opinions, interpretations, conclusions, and recommendations contained herein are those of the authors and are not necessarily endorsed by the U.S. Army.
- The views expressed here are those of the authors and do not necessarily represent the views or official position of the FDA, NIH or DOE.

## AUTHOR CONTRIBUTIONS

H.S. conceived of the project, led the project, collected samples, registered samples, wrote and revised the manuscript, generated figures and tables, performed data analysis, and served as the principal investigator. T.D.M. led the coordination of the use cases and wrote and revised the manuscript. Y.Y. did script/command development, data analysis, gathered and organized FDA-ARGOS database metrics. A.H. performed DNA extraction, library preparation, MIPS and Illumina sequencing, and gathered and organized data for the Ebola in *silico* study, C.S collected and isolated samples, extracted DNA, performed library preparations and Illumina sequencing, gathered and organized data for the *E. avium* gap study, and generated and revised figures, L.T. did IGS sequencing work, L.S. did IGS sequencing work, S.N. did data analysis, and gathered, registered in NCBI and organized data from IGS sequencing work, W.K. and M.S. helped with coordination of BioProject and data submissions and bacterial annotations and the assessment for gap filling, E.H. did viral annotations, D.L. did LMAT analysis, J.A. coordinated LMAT analysis,, J.K. collected and sequenced samples, T.S. helped develop the study and experimental design, S.L. helped develop the study and experimental design, R.S. collected clinical samples, U.S. helped develop the study and experimental design.

