## Supplementary material for "FDA-ARGOS: A Public Quality-Controlled Genome Database Resource for Infectious Disease Sequencing Diagnostics and Regulatory Science Research"

**Supplemental Materials**

**Supplemental Tables (See Attached Excel file):**

1. FDA-ARGOS Reference Genome Database: NCBI Accessions and Assembly Quality Metrics
2. FDA-ARGOS Reference Genome Database: Meta Data Coverage
3. List of Enterococcus avium NCBI Nt Genomes
4. Metagenomics Shotgun Data of Mock Clinical Human Blood Sample Spiked with 10^5^ *E avium*
5. Isolate Shotgun Data of Spiked *E avium*
6. Benchmark and *In Silico* Performance of MIPS BDBV and EBOV Assay
7. CLC Genomics Workbench Analysis of Mock Clinical Human Blood Sample Spiked with 10^5^ *E avium*
8. Limit of Detection (LOD) Table EBOV Serum

**Reference Data Sets :** 5 reference data sets from the use cases are available from Bioproject ID# PRJNA495928 and at <https://www.ncbi.nlm.nih.gov/bioproject/495928>.

1. Metagenomic Shotgun Sequencing for Identification of *E avium*
   a. 3 replicate samples
2. Isolate Shotgun Sequencing for Identification of *E avium*
   a. 3 replicate samples
3. MIPS for Identification of Bundibugyo Virus
   a. 10 PCR positive
   b. 1 NTC
4. MIPS for Identification of Ebola Virus Makona
   a. 15 PCR positive
   b. 1 NTC
5. MIPS EBOV Mock Clinical Trial
   a. 148 blinded samples
      i. 48 positive
         1. 16 10X
         2. 16 5X
         3. 16 1X
      ii. 100 negative (matrix only)

**Additional Files:**

1. FDA-ARGOS Wanted Organism List: Organisms Not Needed (List 1)
2. FDA-ARGOS Wanted Organism List: Priority Organisms (List 2)
3. Recommended Requirements for Nucleic Acid Extractions

**FDA-ARGOS Wanted Organism List**

**List 1: Organisms Not Needed (already covered in FDA-ARGOS with 5 or more genomes):**

**Bacteria**

- ***Achromobacter	xylosoxidans***
- ***Acinetobacter	baumannii***
- ***Alcaligenes	xylosoxidans***
- ***Bacillus	anthracis*, cereus, thuringiensis***
- ***Bordetella	bronchioseptica***
- ***Brevibacterium	casei***
- ***Burkholderia	cenocepacia, cepacia, gladioli, mallei*, multivorans, pseudomallei*, thailandensis***
- ***Campylobacter	jejuni***
- ***Chlamydia	trachomatis***
- ***Chryseobacterium	indologenes***
- ***Citrobacter	freundii, koseri***
- ***Clostridium	botulinum, perfringens***
- ***Corynebacterium	amycolatum, tuberculostearicum***
- ***Delftia	acidovorans***
- ***Enterobacter	aerogenes, cloacae***
- ***Enterococcus	avium, casseliflavus, faecalis, faecium, gallinarum, hirae***
- ***Escherichia	coli***
- ***Francisella	tularensis****
- ***Klebsiella	oxytoca, pneumoniae***
- ***Lactococcus	garvieae, lactis***
- ***Listeria	monocytogenes***
- ***Mycobacterium	tuberculosis***
- ***Neisseria	meningitidis***
- ***Pasteurella	multocida***
- ***Pediococcus	pentosaceus***
- ***Proteus	mirabilis***
- ***Providencia	stuartii***
- ***Pseudomonas	aeruginosa, putida, stutzeri***
- ***Rothia	mucilaginosa***
- ***Salmonella	enterica***
- ***Serratia	marcescens, plymuthica***
- ***Shigella	flexneri, sonnei***
- ***Sphingobacterium	multivorum***
- ***Staphylococcus	aureus, carnosus, hominis, lugdenensis, pasteuri, saprophyticus***
- ***Stenotrophomonas	maltophilia***
- ***Streptococcus	agalactiae, oralis, pyogenes***
- ***Vibrio	alginolyticus, harveyi, parahaemolyticus, vulnificus***
- ***Yersinia	enterocolitica, pestis*, pseudotuberculosis***

**Viruses**

- ***West Nile virus***
- ***Dengue virus***
- ***Venezuelan equine encephalitis virus****

**List 2: Priority Organisms**

**Biothreats (any)**

- ***Ebolavirus****
- ***Crimean-Congo hemorrhagic fever orthonairovirus****
- ***Eastern equine encephalitis virus****
- ***Hantaan orthohantavirus****
- ***Lassa mammarenavirus****
- ***Marburg marburgvirus****
- ***Rift Valley fever virus****
- ***Variola virus****
- ***Western equine encephalitis virus****
- ***Brucella****
- ***Coxiella burnetii****
- ***Rickettsia prowazekii****

**Bacteria (need 1 isolate)**

- ***Acinetobacter	johnsonii***
- ***Acinetobacter	lwoffi***
- ***Candida	parapsilosis***
- ***Citrobacter	amalonaticus***
- ***Clostridium	difficile***
- ***Corynebacterium	glucuronolyticum***
- ***Corynebacterium	striatum***
- ***Granulicatella	adiacens***
- ***Moraxella	catarrhalis***
- ***Moraxella	osloensis***
- ***Morganella	morganii***
- ***Neisseria	gonorrhoeae***
- ***Pediococcus	acidilactici***
- ***Staphylococcus	epidermidis***
- ***Staphylococcus	equorum***
- ***Vibrio	cholera***

**Viruses (any)**

- ***Zika virus***
- ***Chikungunya virus***
- ***Dengue virus***
- ***Ross River Valley Virus***
- ***West Nile virus***
- ***Monkeypox virus***
- ***Chapare virus***
- ***Guanarito Virus***
- ***Hendra virus***
- ***Human coronavirus***
- ***Junin virus***
- ***Lujo virus***
- ***Machupo virus***
- ***Nipah virus***
- ***Rift Valley Fever virus***
- ***Sabia Virus***
- ***Yellow Fever Virus***

**Recommended Requirements for Nucleic Acid Extractions:**

- No isolates or cells will be sent
- The extractions shall be in the following amounts:
  - Bacteria extractions: >15ug (micrograms)
  - Virus extractions: 30ng
  - Large genome extractions: 20-30ug (micrograms)
- Recipient investigator means Dr. Heike Sichtig, PhD
